# *scribble* abrogation induces tumor growth through JNK-Wnt pathways depending on cellular-microenvironment in *Drosophila* wing imaginal tissue

**DOI:** 10.1101/721407

**Authors:** Amarish Kumar Yadav, Roshan Fatima, Saripella Srikrishna

**Affiliations:** Cancer and Neurobiology Laboratory, Department of Biochemistry, Institute of Science, Banaras Hindu University, Varanasi -221005, India; National Centre for Biological Sciences, Bangalore -560065, India

**Keywords:** *scribble*, Cancer, JNK and Wnt signalling, Microenvironment, *Drosophila*

## Abstract

*scribble* (*scrib*) is a cell-polarity determinant in *Drosophila* and human. Cell polarity plays a crucial role in the maintenance of tissue homeostasis and its disruption leads to neoplastic cancer progression. However, the underlying mechanisms by which loss of cell-polarity regulators drives cancer progression are poorly known. In this study, we have explored the tumor progression mechanisms upon *scrib* knockdown in *Drosophila* wing imaginal disc using UAS^RNAi^-GAL4 approach. We have found that *scrib* knockdown in wing disc leads to tumor growth with disrupted actin cytoskeleton, loss of cell-polarity and elevated JNK signalling, resulting in absolute early pupal lethality. Further, *scrib* abrogated cells in a large area of the disc are capable of invading the surrounding wild type cells and inducing apoptosis along with compensatory proliferation through JNK-Wnt pathways. Moreover, JNK pathway upstream candidate *hep* (JNKK) knockdown in *scrib* abrogated cells rescues the cell polarity defects, actin cytoskeleton disruption and tumor growth, while constitutive *hep* activation further aggravates the tumor phenotype. Interestingly, generation of undead cells by apoptosis inhibition in these *hep* knockdown cells by *p35* expression further leads to tumor development. Hence, we conclude that *scrib* knockdown in a large area of wing disc might have a ‘group-protection’ and ‘undead-cells’ microenvironment that modulates the action of JNK signalling resulting in tumor formation. Furthermore, JNK dependent activation of Wnt/Ca^2+^ signalling also supports the tumor growth and actin cytoskeleton disruption. Thus, our results importantly highlight the role of JNK signalling in tumor progression upon *scrib* loss of function depending on cellular-microenvironment.

## 1. Introduction

Major forms of human cancer are epithelial in origin including cervical cancer, breast cancer, prostate and colorectal cancer etc (Pearson et al., 2011; Royer et al., 2011). Proper epithelial cell-polarity is crucial for tissue homeostasis and normal development. Loss of apical-basal cell polarity is one of the hallmarks of the cancerous cell (Ellenbroek et al., 2012; Grifoni et al., 2013). Epithelial cell polarity regulators have been categorised into three major classes known as-Crumbs complex [Crumbs-Pals1 (stardust)-Patj-Lin7] & Par-complex [Par3 (bazooka)-Par6-aPKC] both of which are apically localized and Scrib-Complex (Lgl-Scribble-Dlg), a basolateral determinant and localized towards the basement membrane (Humbert et al., 2006; Royer and Lu, 2011). Among these classes Scrib complex members like Scribble, Lgl & Dlg are well studied in *Drosophila* and reported as neoplastic tumor suppressors (Bilder, 2004).

The *scrib* mutation in *Drosophila* was first isolated by Bilder and Perrimon (2000) in a screen for maternal effect mutations that affect the epithelial morphogenesis. It was named so because of the indented appearance of the embryonic cuticle of the mutants, unlike the smooth surface of wild type. They also cloned the gene and generated antibodies against the protein and showed that it is associated with cell membranes and localizes to the septate junctions. A series of EMS alleles of *scribble* with varying degree of severity have been described (Bilder and Perrimon 2000; Zeitlet et al 2004). While some of them act as null alleles with extended larval life and die as giant larvae, some others act as hypomorphs and die as early to late pupae. Moreover, alleles in certain trans-heterozygous combinations produce escapee flies as well. Apical proteins like armadillo, Sas (Stranded at second), Neurexin, Crumbs, etc. are mis-localized in *scrib* mutants causing disruption of epidermal architecture with groups of cells that are rounded or irregular in shape, a shift from the columnar, mono-layered profile of wild type cells.

Zeitler et al (2004) examined the wing discs of *scrib* mutant larvae using Phalloidin staining that labels the filamentous actin. Wing discs consist of a monolayer of apically elongated epithelial cells with the filamentous actin enriched at the apical end in wild type. However, in *scrib* mutants, actin shows uniform cortical distribution in multi layered groups of cells. The mutant wing discs are larger in comparison to wild type and the phenotype is correlated to the severity of the mutation. Wing discs of null alleles have a higher area, volume and number of cells than those of weaker alleles. Mutations in two other genes, discs-large (dlg) and lethal giant larvae (lgl) also result in loss of apico-basal polarity and over proliferative phenotypes, due to which these genes, along with *scribble*, are known as neoplastic tumor suppressor genes (Brumby and Richardson 2003; Pagliarini and Xu 2003). Moreover, *scrib* is one of the major targets of human papilloma viruses (HPV) and associated with many forms of human-cancer development (Barmchi et al., 2016; Kranjec et al., 2016).

Previous studies in *Drosophila* have shown that small clones of *scrib* mutant do not grow vigorously and are ultimately removed by surrounding wild type cells through cell-competition (Chen et al., 2011), whereas the *scrib* loss-of-function in relatively bigger area resists the removal by wild type cells (Verghese et al., 2012). Further, Chen et al 2011 reported that removal of the *scrib* mutant clones was due to elevated JNK signalling (Chen et al., 2011). However, it has been also shown that active JNK cooperates over proliferation of *scrib* mutant clones with *Ras* gain-of-function (Brumby et al., 2003; Enomoto et al., 2015). Hence, the cellular microenvironment modulates the action of JNK signalling either to promote or suppress the tumor growth. Likewise, *scrib* knockdown in large area shows resistance to its apoptotic removal by surrounding wild type cells suggesting the role of cellular-microenvironment in JNK signalling regulation. But, how large area of *scrib* knockdown cells evade cell-competition mediated removal is still not clear and largely unexplored.

A recent study reported that highly populated *Rab5* mutant cells are able to evade cell-competition through a mechanism of ‘group-protection’ against surrounding wild type cells (Saavedra et al., 2014). These group-protected *Rab5* mutant cells exhibit active JNK signalling and Wg (wingless or Wnt) pathway, supporting overgrowth of wing discs (Saavedra et al., 2014). Wg pathway activation is well reported in many forms of human cancers (Zhan et al., 2017). Wg signaling acts as both canonical (Wg/β-Catenin) and non-canonical (Wg/Ca^2+^) manner to regulate cell proliferation and development (De *et al.*, 2011). Also, it has been reported that in *Drosophila* wing imaginal disc JNK signalling induces Wg and Yki (Yorkie) expression during compensatory proliferation against tissue damage/X-ray irradiation/genetically induced apoptosis (Sun et al., 2011). while, p35 mediated apoptosis inhibition in the X-ray or genetically induced apoptotic-cells, make them ‘undead cell’ which secretes wg/dpp morphogen causing overgrowth of the tissue (Ryoo et al., 2004; Perez-garijo et al., 2009). Wg is an effector mediator of cell competition and up-regulated Wg expression in the cells protect their apoptotic removal. Interestingly, Cell-competition induced dying cells also secret the cell growth promoting morphogen like Wg/dpp and contribute to compensatory proliferation (Levayer et al., 2013).

Loss of cell polarity regulator, Scrib, has been reported from common epithelial human cancers like breast, cervical, colorectal, skin etc (Feng et al., 2016; Kranjec et al., 2016). But, how loss of Scrib promotes tumor growth is largely unexplored. We undertook the present study with an aim to explore the underlying mechanisms of epithelial cancer progression upon *scrib* loss of function. We report that *scrib* knockdown in the wing disc leads to tumor development with disrupted actin cytoskeleton, loss of cell-polarity and elevated JNK signalling resulting in absolute early pupal lethality.

## 2. Results

### 2.1. *scrib* knockdown in wing imaginal discs leads to tumorous growth eventually resulting in early pupal lethality

With an aim to study the mechanism of cancer progression upon *scrib* loss of function, we knocked down the *scrib* in the wing imaginal disc using UAS^RNAi^-GAL4 approach. The expression pattern of Wing-GAL4 was analysed by crossing it to UAS-GFP and UAS-LacZ responder lines, which showed the GAL4 expression in almost the entire disc, including the wing pouch, notum and hinge regions, with an exception of the margins of the disc, as shown in Fig. S1. Thus it was clear that using this approach, it is possible to induce or silence a gene in almost the entire disc. We then went ahead and knocked down *scrib* in the wing discs by crossing UAS-scrib^RNAi^ responder line to Wing-GAL4 driver line. We found that *scrib* knockdown by this approach (UAS-Scrib^RNAi^/Wing-GAL4) is sufficient to promote massive tumour growth with distorted wing-discs morphology causing absolute pupal lethality (Fig.1 A-F). Wing disc area was measured using LSM-image browser software with overlay-drawing mode and measures. Tumorous disc was about 1.3 times larger in area relative to wild type disc (Fig.1 C). Moreover, we observed that *scrib* knockdown driven tumor bearing third instar larvae were relatively larger in size compared wild type larvae. Though they initiate pupation like wild type, the pupae fail to develop further and show 100% lethality at early stages of development, whereas wild type pupae complete their development normally (Fig.1 D-F). Further, immunostaining revealed a reduction in the levels of Scrib expression in the mutant wing discs as compared to wild type (Fig. S2).

**Figure 1.**
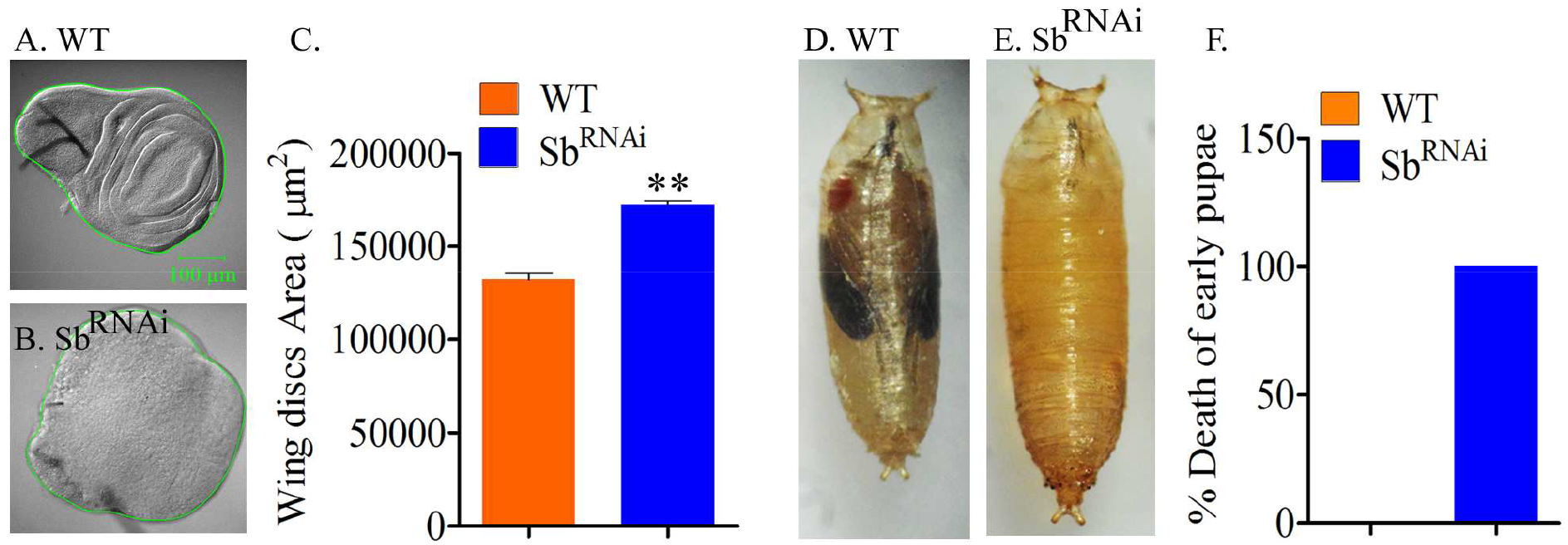
*scrib* Knockdown in wing imaginal discs leads to tumorigenesis and absolute pupal lethality. A-B. DIC images of normal and tumorous wing discs of WT and Sb^RNAi^, respectively. Green circles depict the distorted morphology of Sb^RNAi^ disc as compared to the regular shape of WT disc. **C.** Histogram showing comparative area of WT and Sb^RNAi^ wing imaginal discs. **D-E**. Normally developed pupa of WT and a dead pupa of Sb^RNAi^, respectively. **F**. Histogram showing percent of dead pupae of Sb^RNAi^ (100%, n=129) in comparison to WT (0%).

### 2.2. *scrib* knockdown in wing imaginal discs leads to enhanced JNK signalling

We wanted to investigate the status of JNK signalling in the *scrib* knockdown induced tumor, since it is known that cell-polarity regulators modulate JNK signalling in cancerous tissue (Igaki et al., 2006; Leong et al., 2009). We examined active JNK (Phosphorylated-JNK) in wing imaginal discs as previously shown by Wang et al 2012. *scrib* knockdown induced tumor shows elevated JNK transcription, up to 1.9 folds, compared to wild type disc as revealed by immunostaining and qPCR respectively (Fig.2 A-D and I). Further, we were interested to analyse the presence of proliferating and/or apoptotic cells in this tumor as it is well known that JNK signalling can act as a tumor promoting as well as a tumor suppressing factor. Cell death and cell proliferation were analysed by TUNEL and BrdU assays, respectively in growing tumor against wild type counterpart (Fig.2 E-H). Apoptotic and proliferating cells were counted manually in LSM-image browser software. TUNEL assay revealed that *scrib* abrogated tumor has around 4.6 times higher number of apoptotic cells relative to normal disc (Fig. 2 J), though these apoptotic cells were higher in number towards the edge of tumor (Fig.2 F, white arrows). Moreover, BrdU assay showed the presence of about 1.4 times higher number of actively dividing cells in tumorous disc compared to wild type (Fig. 2 K). However, the actively dividing cells were present throughout the tumorous disc (Fig. 2 H). Also, cyclin-E immunostainig further corroborated the presence of actively dividing cells in growing tumor (Fig. S3). Hence, *scrib* knockdown in the wing disc leads to tumorous growth with elevated JNK signalling resulting in absolute pupal lethality.

**Figure 2.**
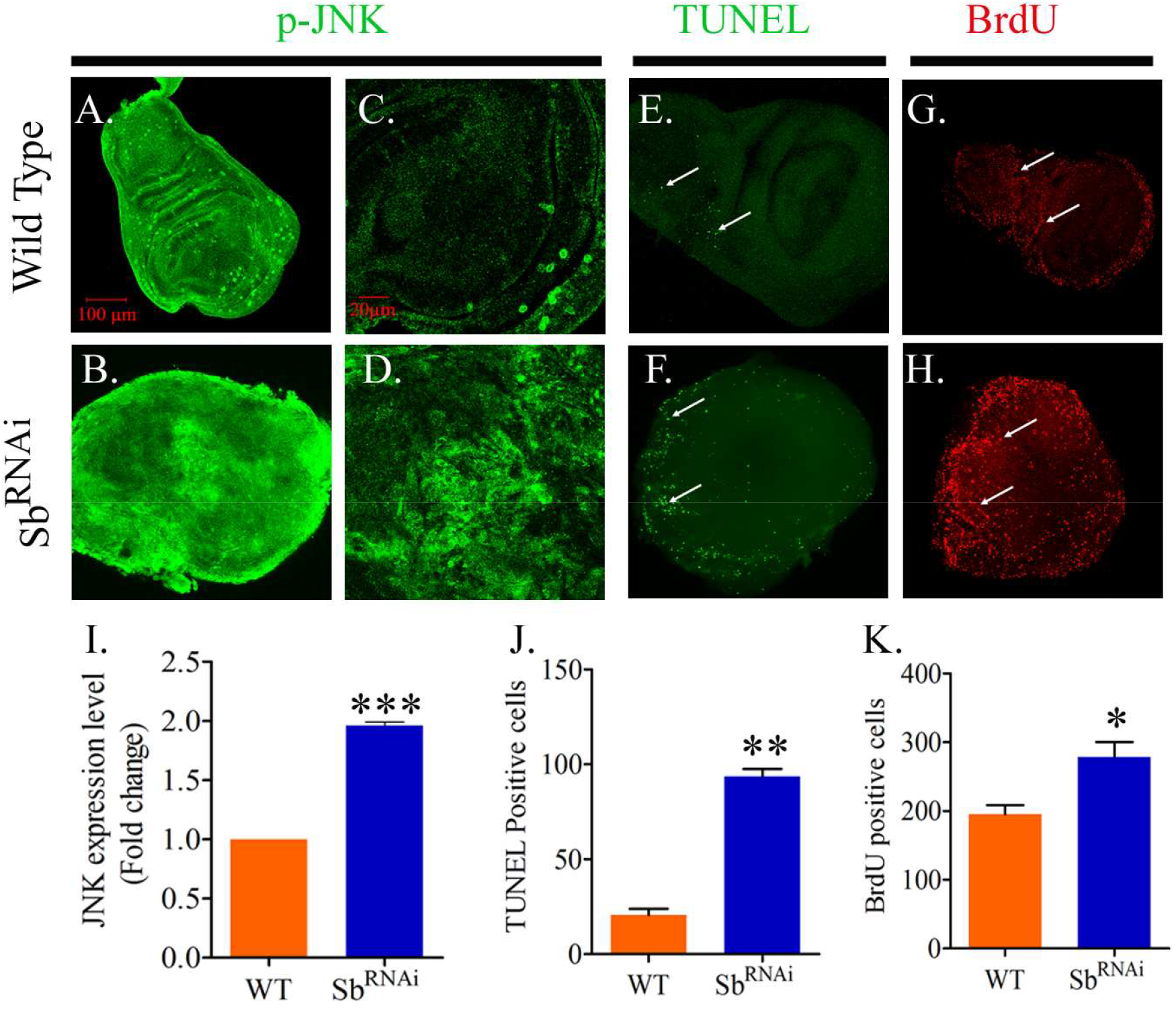
Confocal projections showing active JNK (p-JNK), TUNEL and BrdU immunostaining in Wild type (WT) and *scrib* knockdown (Sb^RNAi^) discs respectively. A-B show wild type (A) and Sb^RNAi^ (B) active JNK (green) status. Sb^RNAi^ shows highly active JNK compared to wild type. Magnified images of the same are shown in C-D. Wild type disc (C) shows normal level of active JNK, though few cells show high level of active JNK (green spots). While,Sb^RNAi^ (D) shows high level of active JNK throughout disc. TUNEL assay (E-F) reveals apoptotic cells (white arrows indicating green dots) in wild type & Sb^RNAi^ discs respectively and Brdu immunostaining (G-H) shows presence of actively dividing cells (red dots indicated by white arrows) in wild type and Sb^RNAi^ discs respectively. Histogram (I) shows about 1.9 folds elevated level of JNK mRNA in Scrib^RNAi^ compared to wild type. (J) and (K) represent the presence of higher number of apoptotic cells and dividing cells in *scrib* knockdown driven tumor compared to wild type counterpart respectively. (p<0.05).

### 2.3. JNK signalling regulates the cell-proliferation and apoptosis in *scrib* knockdown tumor of wing imaginal discs

As we noted that *scrib* knockdown in the wing imaginal disc leads to tumor growth with elevated JNK signalling along with proliferating as well as apoptotic cells, we were interested to understand that how JNK signalling regulates the cell proliferation and cell death in this tumor. Therefore, we genetically modulated the JNK signalling in this tumor by overexpression and knockdown of JNK upstream candidate Hemipterous (Hep) also known as Jun Kinase Kinsae (JNKK). UAS-Hep^Act^ (constitutively active) line was used to up-regulate JNK signalling, while UAS-Hep^RNAi^ line was employed to knockdown the JNK signalling in this tumor. It may be noted that Hep activation causes phosphorylation of Basket (JNK) to induce JNK signalling in the cells (Ages et al., 1999; Stronach, 2005).In accordance with our previous observation, we noted that activation of JNK signalling in the background of *scrib* knockdown induces both cell proliferation as well as cell death, as analysed by BrdU and TUNEL assays, respectively. Activation of JNK signalling in *scrib* knockdown tissues (Wing-Gal4>UAS-Hep^Act^; UAS-Scrib^RNAi^) induces higher cell proliferation, about 8.3% relative to *scrib* knocked down alone (Wing-Gal4>UAS-Scrib^RNAi^) and about 48.5% higher as compared to wild type (Fig.3 A-C, E-G, I-K and Q, Number of Brdu positive cells were counted manually using LSM image browser and calculated in percentage to represent comparisons). We further analysed the cell proliferation in this tumor upon JNK down-regulation (Wing-Gal4>UAS-Hep^RNAi^; UAS-Scrib^RNAi^) and observed that JNK down regulation reduces cell proliferation by about 22.3% relative to *scrib* knockdown alone and also restores the wing disc morphology to near wild type (Fig.3 A-C, E-G, M-O and Q). As far as cell death is concerned, JNK activation in the background of *scrib* knockdown induces higher cell death, about 82.7% as revealed by TUNEL assay (Fig.3 D, H, L and R). We observed that apoptotic cells were present throughout the tumor though more number of apoptotic cells were present towards outer region (Fig. 3L). Likewise, JNK signalling down regulation shows about 13.7% reduction in apoptotic cells in this tumor compared to *scrib* knocked down alone (Fig 3 P and R). Here, we noted that the number of apoptotic cells were still higher in number compared to wild type disc (Fig. 3 R) and were present more towards the inner region of the disc (Fig. 3 P). Hence, JNK signalling modulates the cell proliferation and cell deaths in *scrib* knockdown driven tumor.

**Figure 3.**
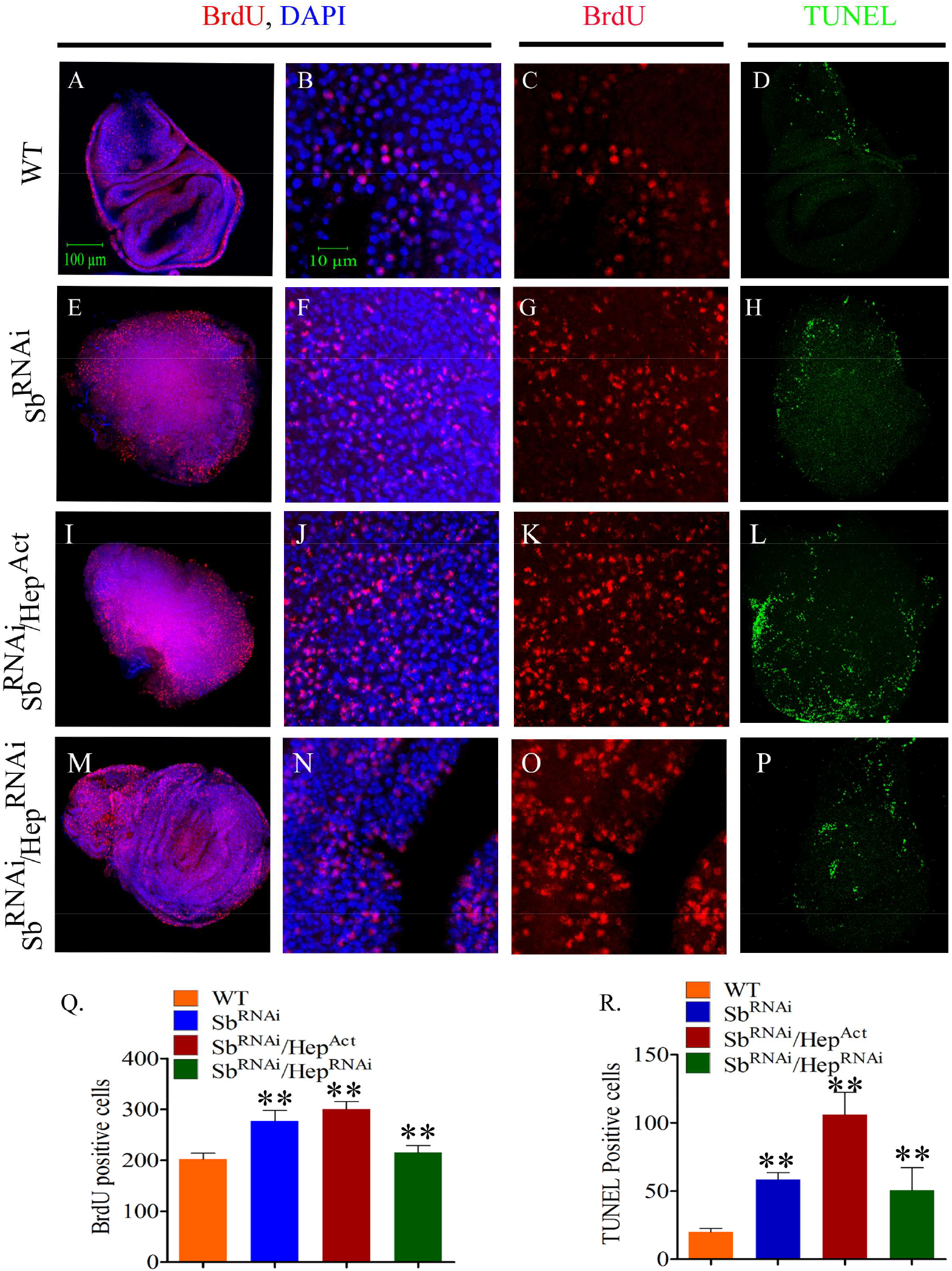
Confocal projections representing BrdU and TUNEL assays. A. Wild type wing imaginal disc showing actively dividing cells (in S-phase) stained with BrdU (red) and DAPI (blue) representing the nucleus. B-C. Magnified wild type disc representing nucleus (blue) and dividing cells (red). Note the nuclear localization of BrdU (B). D. Apoptotic cells (green dots) during wing disc development. Likewise, (E-H), (I-L) and (M-P) show the presence of actively dividing cells and apoptotic cells in *scrib* knockdown upon JNK up and down regulated wing discs respectively. Histograms (Q) and (R) show the number of actively dividing cells and apoptotic cells respectively in corresponding genotypes.

### 2.4. Apoptosis inhibition by *p35* expression in JNK down-regulated *scrib* knockdown disc induces tumorous growth

In previous result, we found that JNK signalling down regulation in *scrib* knockdown cells inhibited the tumor growth but more number of apoptotic cells were present in this disc compared to wild type (Fig.3 D, P). Therefore, we were interested to know what would happen if apoptosis is inhibited in these cells. We used UAS-*p35* line, a caspase inhibitor (Marchal et al., 2012), in these cells to inhibit the apoptosis. We observed that *p35* expression in these cells restores the tumorigenic potential of these cell and induces tumorous growth (Fig.4 A-B). Hence, apoptosis inhibition by *p35* expression in JNK diminished *scrib* abrogated cells further leads to tumor growth.

**Figure 4.**
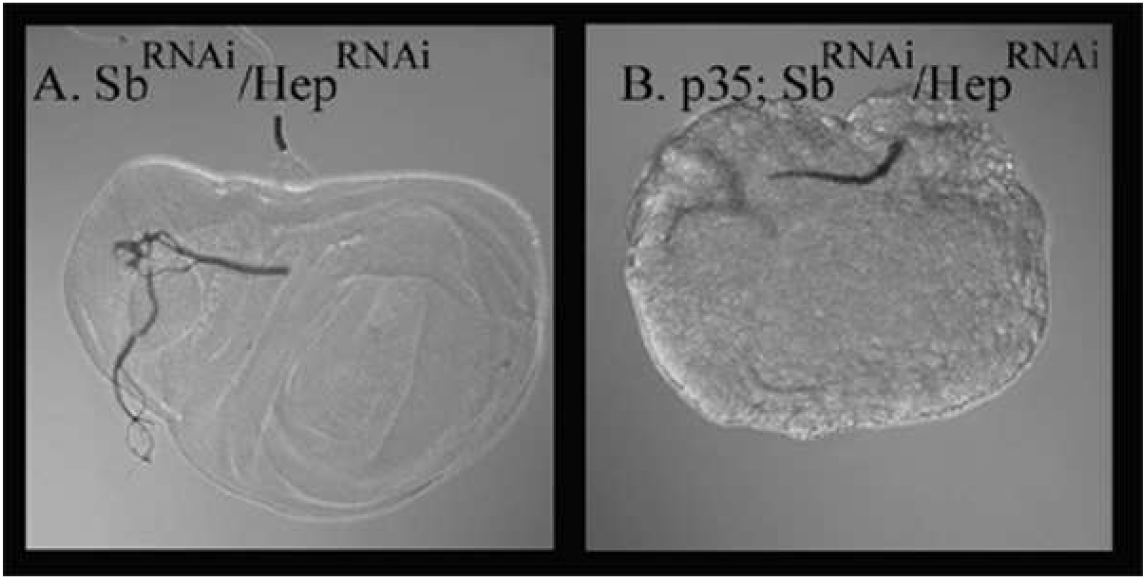
DIC images representing. **A**. JNK down-regulation in *scrib* knockdown cells rescued tumor growth and **B**. *p35* expression in JNK compromised *scrib* knockdown cells induces tumorous growth.

### 2.5. *scrib* knockdown in wing discs show tumorigenesis from early age of larval development and JNK signalling diminution rescues the tumor growth and early pupal lethality

We have observed that *scrib* knockdown tumor grows vigorously with highly disrupted wing disc morphology and elevated level of JNK signalling. So, we were interested to observe how the tumor progresses and what is the fate of *scrib* knockdown cells with JNK signalling up and down regulation, respectively. For this, we marked the cells with *scrib* knockdown alone and in genetic combination of Hep (JNKK), with GFP (green fluorescent protein) using UAS-GFP line and dissected out the wing discs for observation from early to late age of larval development stages i.e. in a window of 36h to 96h after larvae hatched. We found that wild type discs (WingGAL4>UAS-GFP) shows GFP expression present to larger area of the disc from early to late age of development and mostly absent from margins of the disc as shown in merged fluorescence images with DIC images representing whole disc, respectively (Fig5 A-E). While, *scrib* knockdown discs (Wing-Gal4>UAS-GFP; UAS-Scrib^RNAi^) show the distorted discs morphology from early age of development and GFP expressing cells are progressively expanded throughout the discs and at late age of 72-96 hrs these cells covers the whole area of the discs and reach the outer margin as well (Fig.5 F-J). Likewise, JNK signalling up regulation (Wing-Gal4>UAS-Hep^Act^/UAS-GFP; UAS-Scrib^RNAi^) in this tumor further aggravates the disc morphology distortion from early age and GFP expressing cells invade the outer margins to covers the whole disc (Fig.5 K-O). However, JNK down-regulation in *scrib* knockdown discs (Wing-Gal4>UAS-Hep^RNAi^/UAS-GFP; UAS-Scrib^RNAi^) show the progressive removal of GFP expressing *scrib* knockdown cells from the early age of development to restore the normal wing disc morphology (Fig.5 P-T).

**Figure 5.**
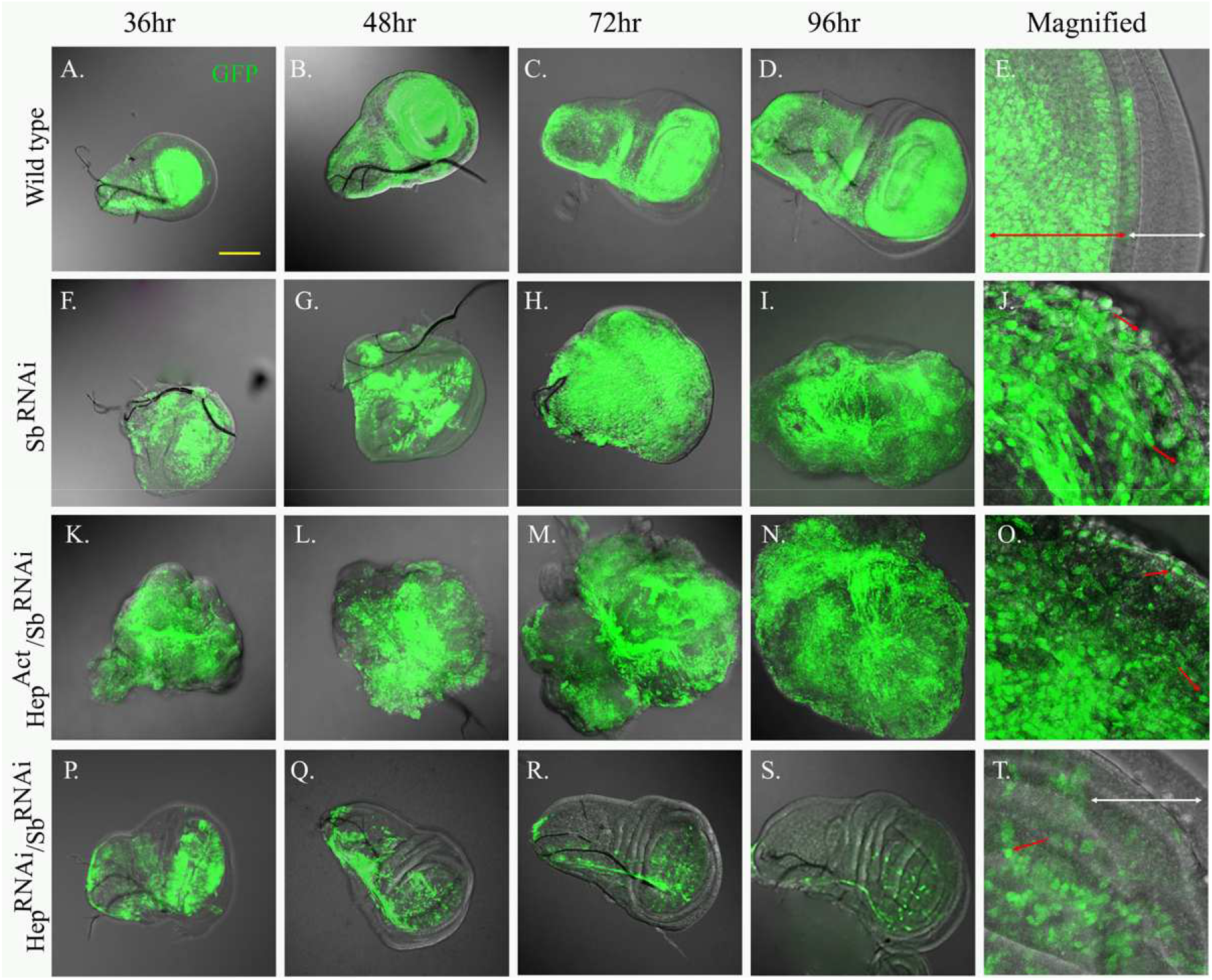
Confocal projections representing GFP (green) expression in developing wild type and tumorous discs at different time frames after larvae hatched. (A-D) wild type shows GFP expression throughout the disc from early to late stages of development except at the margins. (E) Magnified wild type wing pouch region represent GFP expression restricted to pouch region (red arrow) and absent from outer margin (white arrow). Likewise, (F-I) Sb^RNAi^ (WingGal4>UAS-GFP; UAS-Scrib^RNAi^) and (K-N) Hep^Act^/Sb^RNAi^ (Wing-Gal4>UAS-GFP/UAS-HepAct;UAS-Scrib^RNAi^) show GFP expression throughout and late age discs shows invasion of GFP cells at outer margin also. Magnified (J, O) show GFP cell at outer region suggestion invading nature of *scrib* abrogated cell leading to loss of wild type cells, JNK activation further enhances invading capacity of scrib abrogated cells. While (P-S) Hep^RNAi^/Sb^RNAi^ (WingGal4>UAS-GFP/Hep^RNAi^;UAS-Scrib^RNAi^) JNK signalling down regulation shows loss of GFP expressing *scrib* abrogated cells and their replacement by wild type cells. (T) Magnified image shows loss of GFP cells from wing pouch region in comparison to wild type (E) and *scrib* knockdown disc (J) (n=10). (Scale bar represents 100µm).

Moreover, we found that JNK down regulation in *scrib* knockdown wing imaginal discs protect early pupal deaths significantly (82.2%) and the pupae reach upto pharate adult stage (Fig. 6 A, D). Also, some of these puape (7%) were able to eclose as adult flies with distorted (around 40.5%) as well as very normal wings (39.5%) morphology. Distorted wings show the developmental defects in Anterior Cross Vein (ACV) and Posterior Cross Vein (PCV) (Fig. 6B, C, E). Though, eclosed flies show life span near to wild type flies (Fig. 6F). Hence, *scrib* knockdown tumor grows progressively in JNK dependent manner and JNK down regulation protects the tumor formation and associated anomalies.

**Figure 6.**
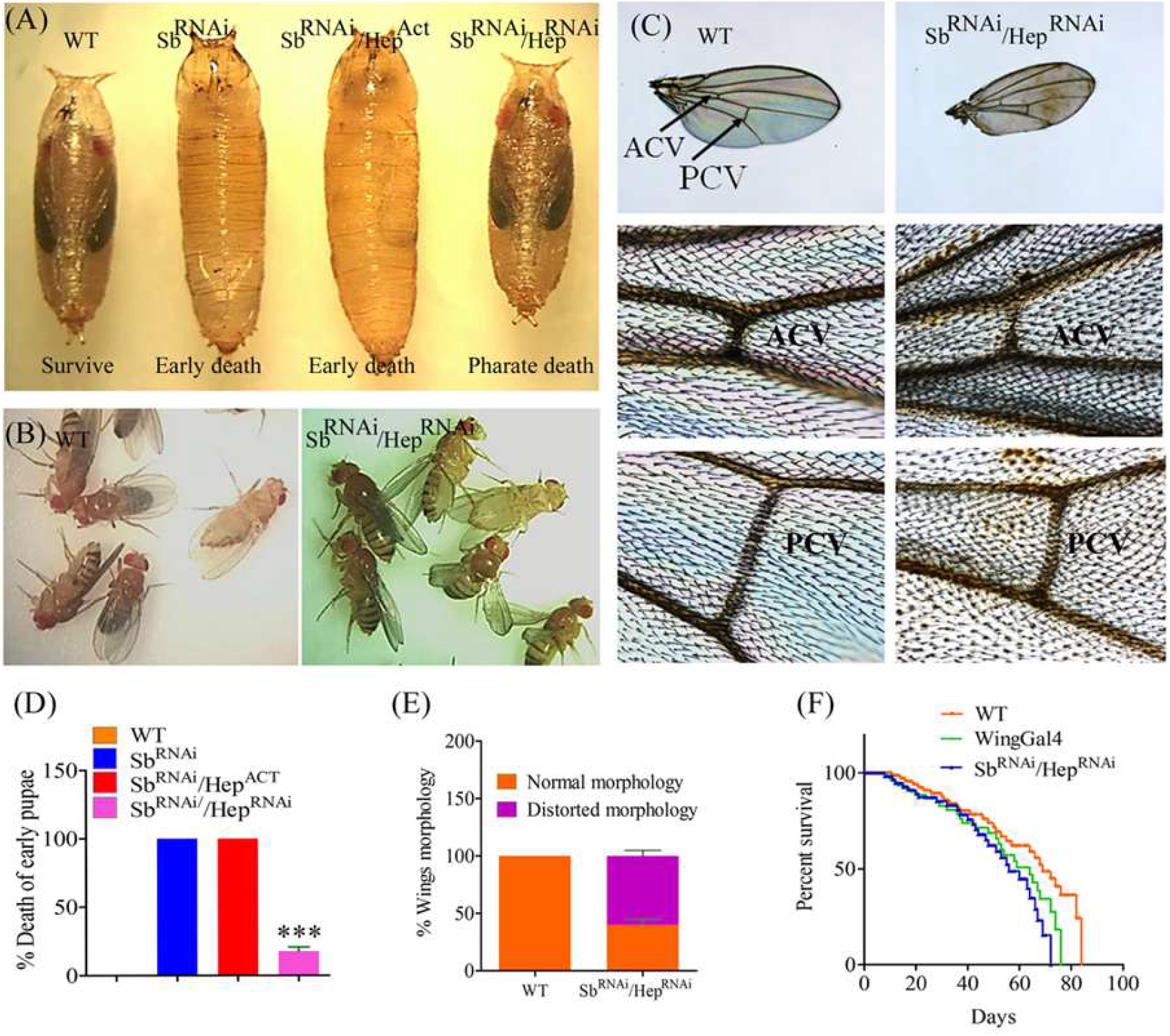
Pupal development, adult wings morphology and percent survival of escaper flies. A. WT (Wild type) shows the fly development inside pupal case while Sb^RNAi^ (WingGal4>UAS-Scrib^RNAi^) and Sb^RNAi^/Hep^ACT^ (WingGal4>UAS-Hep^Act^;UAS-Scrib^RNAi^ lead to early pupal death and enlargement of pupa size, But Sb^RNAi^/Hep^RNAi^(WingGal4>UAS-Hep^RNAi^;UAS-Scrib^RNAi^) showed normal pupal development, although they died as pharate adults, however very few were able to eclose. (B) Wild type (WT) and Sb^RNAi^/Hep^RNAi^ escapers flies. (C) Bright field images showing wing morphology of wild type and escapers flies. Magnified images below show the developmental defects of Anterior Cross Vein (ACV) and Posterior Cross Vein (PCV) of the escaper flieswith relatively smaller size of cross veins. The yellowish-black spots in escapee fly wings might be melatonic depositions. Histograms (D-E) represent percent death of early pupae for respective genotypes (n=115) and wing disc morphological defects (n=20) respectively (***p<0.001). (F) Shows life span of escapee flies compared to wild type and WingGAL4 driver line (n=40).

### 2.6. JNK signalling down-regulation restores actin cytoskeleton and cell polarity in *scrib* abrogated wing imaginal disc

Scrib is one of the cell polarity regulators restricted towards basolateral side of the cell in complex with Dlg and Lgl. So, we analysed the cell polarity in this tumor and actin cytoskeleton rearrangement, a property of tumorous tissue (Efremov et al 2015). Actin cytoskeleton organization was analysed by Phalloidin staining in tumorous disc compared to wild type disc, while cell polarity was observed by basolateral cell polarity marker, Dlg (disc large) immunostaining. We found that actin cytoskeleton was highly disrupted in tumorous disc compared to wild type. Scrib knockdown and JNK activation (WingGAL4>UAS-Hep^Act^; UAS-Scrib^RNAi^) driven tumor shows highly polymerized and accumulated actin filaments, while JNK signalling down-regulation (WingGAL4>UAS-Hep^RNAi^; UAS-Scrib^RNAi^) restores the actin cytoskeleton arrangement comparable to wild type counterpart (Fig.S4 A-H). Likewise, we found that *scrib* knockdown driven tumor cells show loss of basolateral cell polarity as revealed by mislocalized Dlg (appears to be present along the cell-membrane and in cytoplasm) compared to wild type counterpart. Further, we found that JNK signalling activation in this tumor shows Dlg mislocalization though it is similar to *scrib* knockdown tumor, while JNK signalling down regulation restores the cell polarity with restricted distribution of Dlg to basolateral side (Fig. S4, I-T). Hence, restoration of actin rearrangement and cell polarity further corroborate the replacement of *scrib* knockdown cells by wild type cells upon JNK down-regulation as shown in Fig.5S.

### 2.7. *scrib* knockdown driven tumor shows altered levels of Wingless, β-catenin and intracellular Ca^2+^ in parallel to JNK signalling

Since *scrib* knockdown tumor grows with simultaneous presence of apoptotic as well as proliferating cells in JNK dependent manner, we were interested to analyse the Wingless (Wg) pathway in this tumor, as Wg pathway has been shown to be involved in apoptosis induced compensatory proliferation (Ryoo et al 2004). Here, we observed the Wg (also called Wnt) and its downstream effector β-catenin (Armadillo) levels by immunostainings in this tumor in respect to JNK signalling up and down regulation, respectively. *Scrib* abrogated tumor (Wing-Gal4>UAS-Scrib^RNAi^) shows elevated Wingless (Wg) signalling compared to wild type counterpart as revealed by immuno-staining. Also, wild type wing imaginal disc shows a characteristic expression pattern of Wg along the wing-pouch and notum regions, however, it’s hard to identify the wing pouch and notum area for growing tumor (Fig.7 A, B, E, F). Further, JNK signalling up regulation induces the Wg expression, while JNK down regulation reduces the Wg expression in this tumor relatively (Fig. 7 C, D, G, H). Further, β-catenin (Armadillo) level was found to be increased in *scrib* knockdown tumor compared to wild type and it also alters with respect to JNK signalling up and down-regulation as revealed by immunostaining. (Fig S5).

**Figure 7.**
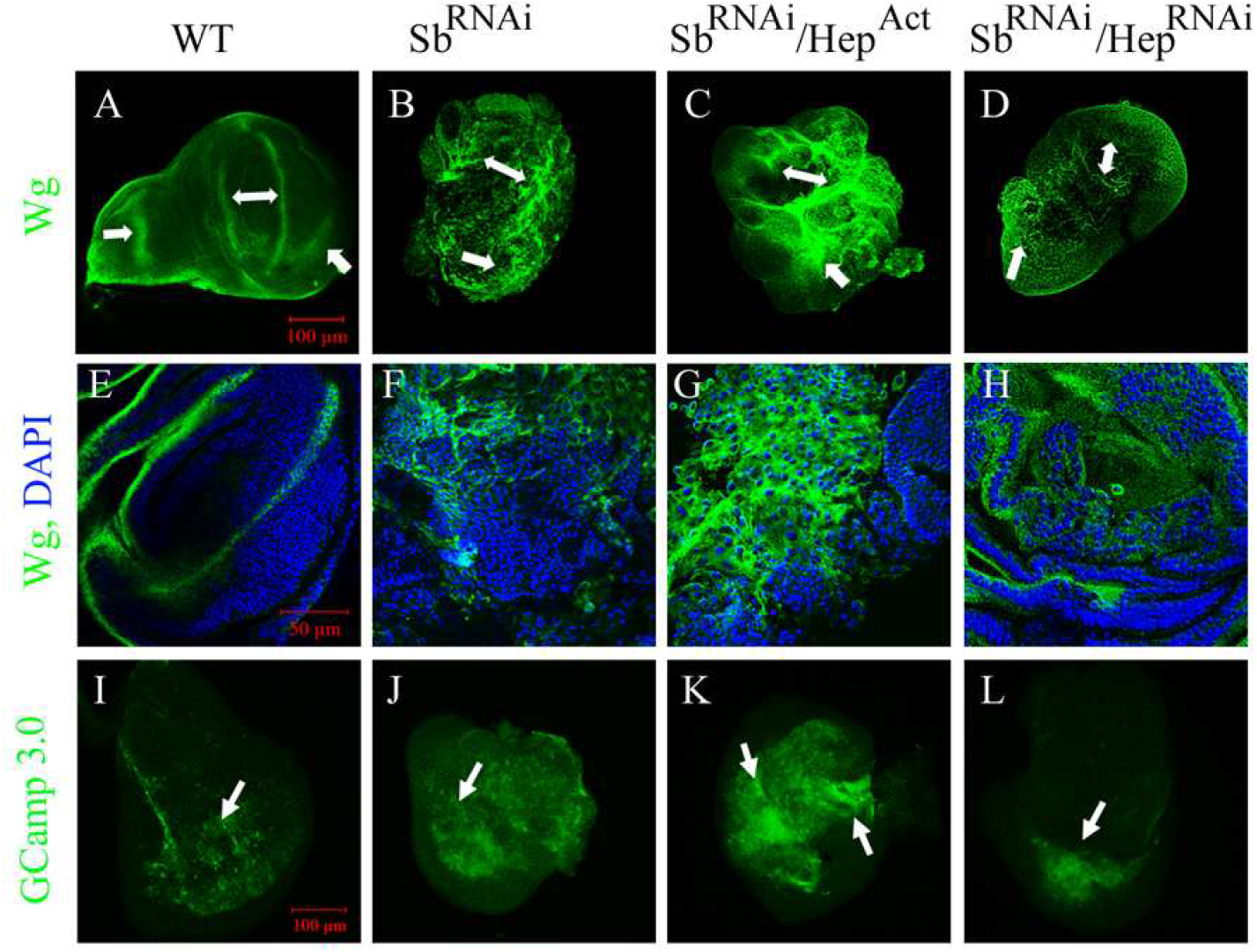
Confocal images of wing imaginal discs representing Wg (wingless) immunostaining and Ca^2+^ detection using GCamp3.0 detector. (A, E) Wild type wing imaginal discs representing wingless (green) pattern (arrows) while (B, F) *scrib* knockdown wing disc tumor shows higher level of Wg expression and (C, G) constitutive Hep (JNKK) activation further enhances Wg expression, But (D, H) decreasing Hep expression reduces Wg expression in *scrib* knockdown tumor. Lower panel (I-L) represent cytosolic Ca^2+^ (green) status in wild type wing disc and tumor tissue. (I) WT shows low cytosolic Ca^2+^ (arrow) in comparison to (J) loss of *scrib* driven tumor while (K) Hep activation further enhances cytosolic Ca^2+^ level and (L) reducing hep activity reduces Ca^2+^ level relatively. (n=10).

Moreover, we analysed the intracellular level of Ca^2+^, another ‘Wnt’ transducer besides β-catenin. Also, Ca^2+^ is known to facilitate the β-catenin entry into nucleus to induce transcription (Thrasivoulou et al, 2013). We observed the Ca^2+^ level in this tumor using UAS-GCamp3.0 responder line, which encodes a calcium indicator that fluoresces green in presence of intracellular Ca^2+^. *scrib* knockdown driven tumor (WingGAl4>UAS-GCamp3.0; UAS-Scrib^RNAi^) shows elevated level of Ca^2+^ relative to control disc (WingGAl4>UAS-GCamp3.0). Further, JNK signalling up-regulation (WingGAl4>UAS-GCamp3.0/UAS-Hep^Act^; UAS-Scrib^RNAi^) enhances the intracellular Ca^2+^ while, JNK signalling down-regulation (WingGAl4>UAS-GCamp3.0/UAS-Hep^RNAi^; UAS-Scrib^RNAi^) reduces the Ca^2+^ level relative to *scrib* knockdown tumor alone (Fig. 7 I-L). Hence, *scrib* knockdown tumor shows altered level of Wg/β-catenin/Ca^2+^ in parallel to JNK signalling.

## 3. Discussion

*Drosophila* larval imaginal discs (e.g. wing imaginal disc) are epithelial monolayer structures, which have been extensively used by researchers to study various aspects of development and disease-biology, including cancer (Froldi et al., 2008). Most of the human cancers are epithelial in origin and are characterised by loss of cell polarity (McCaffrey et al, 2011). Scrib is a well-studied cell polarity regulator and its loss-of-function is associated with various forms of human-epithelial cancer (Kapil et al., 2017). In our study, we have knocked down *scrib* gene in large area of wing imaginal disc to induce tumor growth, using UAS-GAL4 system. We confirmed the expression of Wing-GAL4 (used in this study) to large area of the disc by GFP and LacZ expression analysis (Fig S1). Scrib immunostaining revealed the reduced level of Scrib compared to wild type wing disc (Fig. S2). We reported that *scrib* knockdown (UAS-Scrib^RNAi^/WingGAL4) in large area of wing disc resulted in tumorous growth causing absolute early pupal lethality (Fig. 1 A-F). Also, previous studies with various alleles of *scrib* mutants had shown similar findings with varying degrees of developmental disparity leading to embryonic lethality, wing disc tumor growth and pupal lethality (Bilder et al, 2000 and Zeitler et al. 2004). Scrib knockdown driven tumor of wing imaginal disc might have a systemic effect causing developmental defects associated with the early pupal lethality (Clarevega et al 2015). We have analysed the role of JNK signalling in *scrib* knockdown driven tumor, since previous studies reported the activation of JNK signalling upon loss of cell polarity regulators but the precise role of JNK signalling in these tumors is unknown and remained to be explored. First, we analysed the transcript levels of JNK by qPCR and the status of phosho-JNK (p-JNK) by immunostaining in *scrib* knockdown tissues. We found that *scrib* knockdown leads to transcriptional up-regulation of JNK as well as induction of JNK signalling (Fig 2I). In *Drosophila*, active JNK signalling upon loss of cell-polarity regulator Scrib has been shown to cooperate the oncogenic signals to cause over proliferation in eye imaginal disc (Brumby et al., 2003). However, JNK mediated apoptosis has also been reported in the removal of small *scrib* mutant clones in eye imaginal tissue (Chen et al., 2011), suggesting the context dependent role of JNK signalling. In the current study, we report that active JNK signalling induces tumorous growth upon *scrib* knockdown in large area of wing imaginal disc.

To explore the role of JNK signalling in tumor progression, we examined the cell proliferation and cell death status in *scrib* knockdown driven tumor. BrdU and TUNEL assays revealed the presence of proliferating as well as apoptotic cells in the *scrib* knockdown induced tumor. However, the apoptotic cells were majorly present towards edge of the tumorous disc (Fig. 2 F) and proliferating cells were observed throughout the disc (Fig 2 H). We found that genetically up regulated JNK signalling (through constitutively active hep, UAS-Hep^Act^) further induces the cell proliferation as well as cell deaths in the growing tumor, though again the majority of apoptotic cells were present towards the edge (Fig 3. L). Correspondingly, JNK signalling down regulation by UAS-Hep^RNAi^ in *scrib* knockdown tumor reduces the number of apoptotic and proliferating cells relative to *scrib* knockdown alone, however in this case the apoptotic cells were relatively more towards the wing pouch and notum regions (Fig. 3 P). These observations clearly suggested the critical role of JNK signalling in cell death and cell proliferation determination in *scrib* knockdown induced tumor. Further, we analysed the fate of GFP marked *scrib* knockdown cells upon modulation of JNK signalling. As tumor grows from early to late age, the GFP positive cells progressively spread throughout the disc suggesting invasive nature of *scrib* abrogated cells (Fig5 F-J). Moreover, JNK activation further cooperates with invasive property and leads to a rapid growth of tumor from early age of development (Fig.5 K-O). Also, it is anticipated that presence of GFP expressing cells in outer margin regions of the tumorous disc might be due to competitive removal of wild type cells (Amoyel et al, 2014) as suggested by the presence of apoptotic cells at outer margin area (Fig. 2F). Interestingly, JNK down-regulation in *scrib* knockdown cells (Wing-Gal4>UAS-Hep^RNAi^/UAS-GFP; UAS-Scrib^RNAi^) restore normal wing discs morphology with progressive removal of GFP expressing *scrib* knockdown cells from the disc from an early age of development (Fig.5 P-T). Removal of GFP expressing *scrib* knockdown cells from the disc suggested the cell-competition mediated removal of JNK compromised *scrib* knockdown cells by wild type cells and restoring normal wing disc morphology. Also, presence of apoptotic cells towards wing pouch and notum region further strengthen the removal of *scrib* knockdown cells by neighbouring wild type cells (Fig. 3P). Progressive loss of GFP expressing *scrib* knockdown cells of wing disc show the cell-competition at interface of wild type cells. Though, even at late stage presence of GFP expressing cells in the core of wing pouch suggested that inner cells not facing direct cell-competition are protected and must be slow growing. Wild type cells in the margins were capable of removing the JNK compromised *scrib* abrogated cells to revert the wing disc morphology leading to normal pupal development to distinct wing, eye and complete fly development (Fig. 6). Hence, our study indicates that *scrib* knockdown cells in a large area were group-protected and modulate the role of JNK signalling in cancer progression.

Parallelly, we suggest that the tumor growth upon p35 inhibited apoptosis in JNK diminished *scrib* abrogated cells (Fig. 4) must be due to the ‘undead-cell’ behaviour of inner cells not facing cell competition. These inner undead-cells might be supporting the survival of *scrib* knockdown cells at wild type interface leading to over-proliferation. It has been reported that undead-cells secret cell-growth promoting morphogenes like wg/dpp etc causing over-proliferation (Huh JR et al; 2004). Also, wingless up-regulation along with JNK in group-protected *Rab5* mutants is known to supports such kind of cell-proliferation in wing disc (Arias et al; 2014). Hence, the elevated level of wingless (Wnt) and armadillo (β-catenin) in *scrib* abrogated tumor might be a suggestive cause of cell proliferation even with JNK activation (Fig 7). Further, altered wingless expression in *lgl* mutant (Mukherjee and Roy, 1995) has indicated the role of wingless in tumor progression upon loss of cell polarity regulators. Also, the wingless up-regulation is required in mutant cells to make them ‘super-competitor’ to protect their removal by neighbouring wild type cells through cell-competition (Tamori et al, 2011; Amoyel et al, 2014). These super-competitor cells with enhanced wg expression induce apoptosis in nearby wild type cells, consequently the dying cells also release morphogenes like wingless supporting compensatory cell-proliferation (Amoyel et al 2014; Claver and Torres, 2016). In *scrib* abrogated discs, the presence of higher number of actively dividing cells in close vicinity of apoptotic cells at interface of wild type cells suggested wingless mediated compensatory proliferation (Fig S6).

In addition, the previous report by Zhu et al have proposed the role of JNK signalling in cell polarity regulator *lgl* mutant cells to regulate cell-proliferation, cell-polarity and actin cytoskeleton dynamics (Zhu et al 2010). Likewise, we have found that *scrib* knockdown cells with active JNK signalling show the disrupted cell-polarity and actin cytoskeleton arrangements (Fig S5). The suggestive cause of JNK mediated actin remodelling and cell polarity disruption might be due to activation of Wnt/β-catenin/Ca^2+^ signalling pathways (Tejada-romero et al., 2015; Zhan et al., 2017). Moreover, the JNK mediated transcriptional regulation of Wnt has been reported in eye disc to modulate cell deaths (Zhang et al., 2015; Tare et al., 2016). Further, we have analysed that JNK activation induces Wnt (Wingless) expression in *scrib* abrogated cells along with increased intracellular Ca^2+^ level in cytoplasm (Fig 7. I-L). Wnt expression is known to elevate cytoplasmic Ca^2+^ to facilitate β-catenin entry to the nucleus and also induces non-canonical Wnt/Ca^2+^signalling which regulates the cell-polarity and actin dynamics (Thrasivoulou et al., 2013; Zhan et al., 2017). Hence, our data suggested that in *scrib* abrogated wing imaginal disc JNK mediated Wnt non-canonical signalling must be influencing cell-polarity and actin cytoskeleton arrangement.

In conclusion, our findings highlight the fact that *scrib* knockdown induced JNK signalling disrupts actin dynamics and induces cell proliferation through activation of Wnt pathway. More importantly, the cellular microenvironment like ‘group protection’ and ‘undead cells’ behaviour modulates the action of JNK signalling to augment carcinogenesis (Fig8) in large area *scrib* abrogated cells.

**Figure 8.**
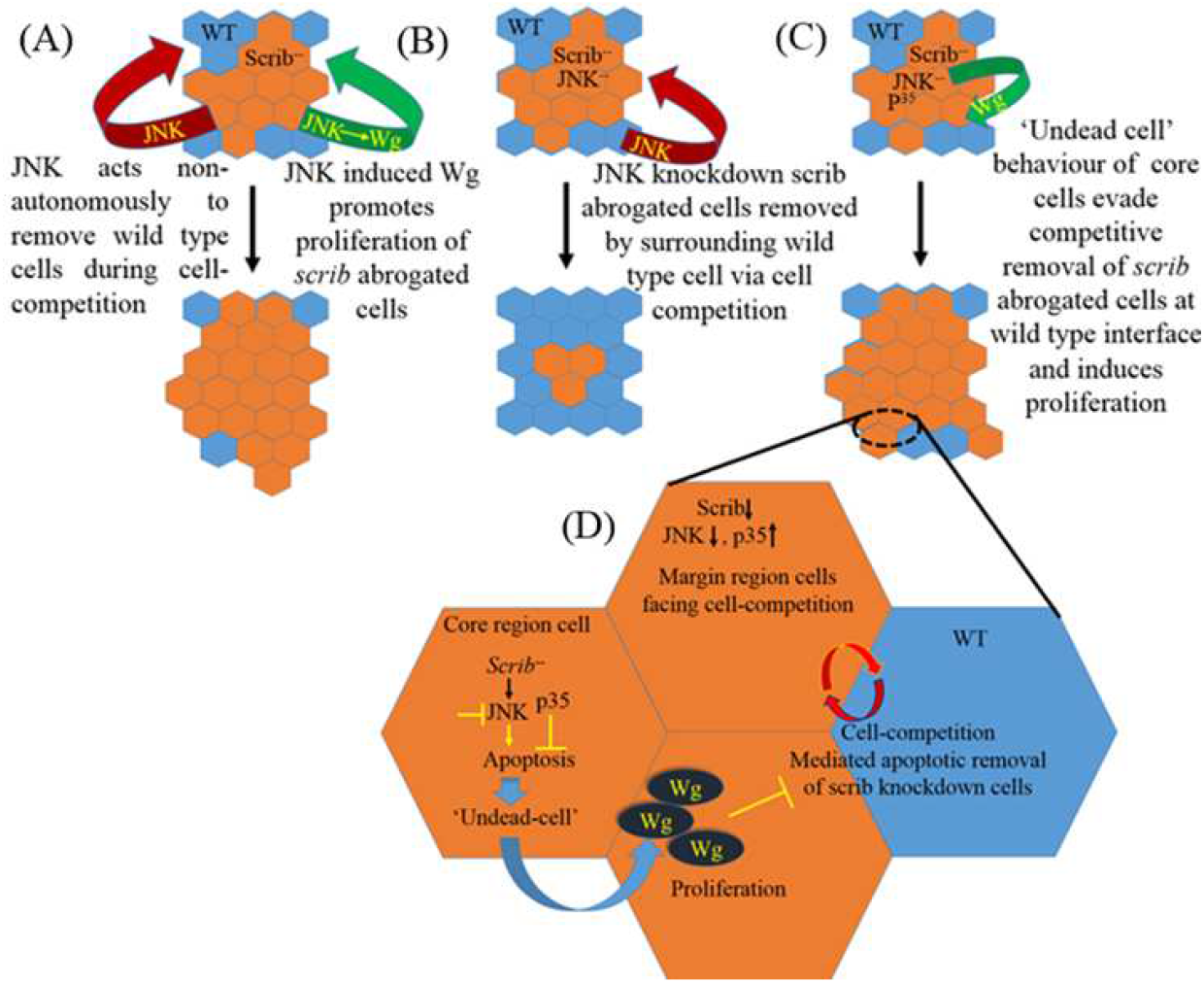
Schematic diagram representing cell-competition and ‘undead cell’ mediated tumor growth upon *scrib* abrogation in large area of wing imaginal disc. (A) Shows active JNK pathway in *scrib* knockdown cells acts non-autonomously to remove surrounding wild type cells through cell-competition and active JNK induces cell proliferation by inducing wg signalling, (B) shows competitive removal of JNK down regulated *scrib* knockdown cells by wild type cells to restore the tissue morphology while (C) represents apoptosis inhibition by overexpression of *p35* in JNK compromised *scrib* knockdown cells interestingly restores the proliferation potential of the *scrib* abrogated cells. In set of lower projected cells (D) represent a proposed mechanism protecting *scrib* and JNK knockdown cells from cell-competition mediated removal by neighbouring wild type cells and leading to over proliferation as ‘undead’ core cells supporting the survival of marginal cells through wg signaling making them super-competitor.

## 4. Material and methods

### 4.1. Fly stocks and culture conditions

Wild type (Oregon-R), UAS-*scrib*^*RNAi*^(#35748),UAS-GFP (#1521), UAS-LacZ, UAS-Hep^ACT^(#9306), UAS-Hep^RNAi^ (#2190-R2), (UAS-GCamp3.0); DM/TM3sb, Sp/Cyo; dCO2/TM6B and Wing-GAL4 (#32120) stocks used in study, were procured from Bloomington *Drosophila* Stock Center, USA. *Drosophila* flies were cultured on standard cornmeal agar media and stocks were maintained in BOD with alternative light-dark cycle at 24±1°C.

### 4.2. X-Gal staining

Third instar larvae were dissected in 1X PBS to isolate wing imaginal discs in cavity slides followed by washing with 50mM phosphate buffer (pH 8) then fixed in 2.5% gluteraldehyde for 10 min each at room temperature. Fixative was removed and tissues were washed with wash buffer (50mM PB) three times each, then X-gal solution was added and kept in dark at 37°C for 30 min followed by washing with wash buffer to remove staining solution. Imaginal discs were transferred onto fresh slides and mounted in 50% glycerol, cover slips sealed with DPX to observe under compound microscope.

### 4.3. Immunohistochemistry

Third instar larvae were dissected in 1X PBS to separate wing imaginal discs and then fixed in 4% paraformaldehyde in PBS for 20 minutes at room temperature followed by washing with PBST (1X PBS and 0.1% Triton X-100), 3 times for 10 min each. Then wing imaginal discs were incubated in the blocking buffer containing 3% bovine serum albumin in PBS 0.3% Tween-20 for 1hr at room temperature. Tissues were incubated with primary antibody in blocking solution for overnight followed by washing with PBST, 3 times for 15 min each. Then the tissues were incubated with secondary antibody in 1XPBS for 3 hr at room temperature followed by washing with PBST, 3 times for 10 min each. These tissues were further incubated with Phalloidin (1:50) for 45 min at room temperature followed by washing with PBST for 3 times 5 min each and then incubated with DAPI (1µ g/mL in 1X PBS) for 10 min, washed in 1X PBS 3 times for 5 min each and subsequently mounted on glass slides in DABCO. Images were obtained on confocal microscope (Zeiss LSM-510 Meta). Primary antibodies used were: goat anti-scrib (Santa Cruz) at 1:50, mouse anti-Wnt (DSHB) at 1:30, mouse anti-Dlg (DSHB) at 1:30, mouse anti-BrdU (DSHB) at 1:20, rabbit anti-pJNK (Promega) at 1:100, and Secondary antibodies were anti-goat-IgG-TRITC (1:200), anti-mouse-IgG-FITC (1:200) and anti-mouse-IgG-TRITC (1:200) were used for respective experiments.

### 4.4. BrdU staining

Third instar larvae were dissected in 1XPBS and wing imaginal discs were isolated, incubated in BrdU solution (0.01mg/ml in 1XPBS) for 1hr at RT followed by 3 times washing with 0.1% PBST for 5min each then fixed with 4% paraformaldehyde for 20 min and followed by 3 times washing with PBST for 5 min each, then treated with 2N HCl for 20 min and washed 3 times with PBST for 5min each followed by blocking in blocking solution for 1hr at room temperature then incubated with anti-BrdU primary antibody (mouse-anti-BrdU, 1:100) for overnight and washing with PBST for 3 times 5 min each and then incubated with secondary antibody for 3 hr and washing with PBST for 3 times 5 min each and then mounted wing discs in DABCO solution to observe under confocal microscope.

### 4.5. TUNEL assay

Third instar larvae were dissected to isolate wing imaginal discs in 1XPBS followed by fixation with 4% paraformaldehyde for 15min and washed for 3 times with 1X PBS followed by permeabilization with 0.1% Triton-X in 1XPBS for 5min and washed three times with 1X PBS 5 min each followed by incubation with TUNEL Assay reaction mixture at 37°C for 2 hr as per manufacturer’s instruction followed by washing with 1XPBS for two times 5 min each and then discs were mounted in DABCO to observe under confocal microscope.

### 4.6. Real time PCR

Total RNA was isolated from 60 wing imaginal discs each from 3rd instar larvae of respective genotypes using TRI reagent (TAKARA) as per manufacturer’s protocol. High quality RNA from both the samples (as estimated by absorbance ratio A260/280) was reverse transcribed for cDNA synthesis through reverse transcriptase PCR. cDNA of the respective sample was used as template DNA for quantification of jnk normalised against *gapdh* using specific primer sets of: *jnk*: FP-CACCAACACTACACCGTCGA; RP-AAGCGGCGCATACTATTCCT, *gapdh*: FP-TAAATTCGACTCGACTCACGGT; RP-CTCCACCACATACTCGGCTC by SYBR green JUMPSTART TAQ Ready mix (Thermofisher Scientific) using real time thermal cycler (Applied Biosystems 7500 real time PCR). All samples were run in triplicate and expression levels were determined by calculating (ΔCT values) threshold cycle analysis.

### 4.7. Microscopy and image processing

Samples were analysed by confocal microscopy using LSM-510 Meta microscope. Optical sections were captured using ZEN Black 2009/LSM image browser software followed by processing in Adobe Photoshop^(R)^ 7.0 to assemble the images.

## Acknowledgments

This work was supported by a research grant (P07/598) awarded to S.S by the Science and Engineering Research Board, India and authors are highly grateful to SERB for financial support (SERB_2015-19). The authors would like to greatly acknowledge Cytogenetics Laboratory, Dept. of Zoology for providing Confocal microscope facility, also ISLS-BHU and UPE-BHU for Nanodrop, Real time PCR and other routine facilities. One of the authors (AKY) is grateful to ICMR and DST-PURSE for providing financial support in the form of Senior Research Fellowship.

## Supplementary Information

**Figure S1.**
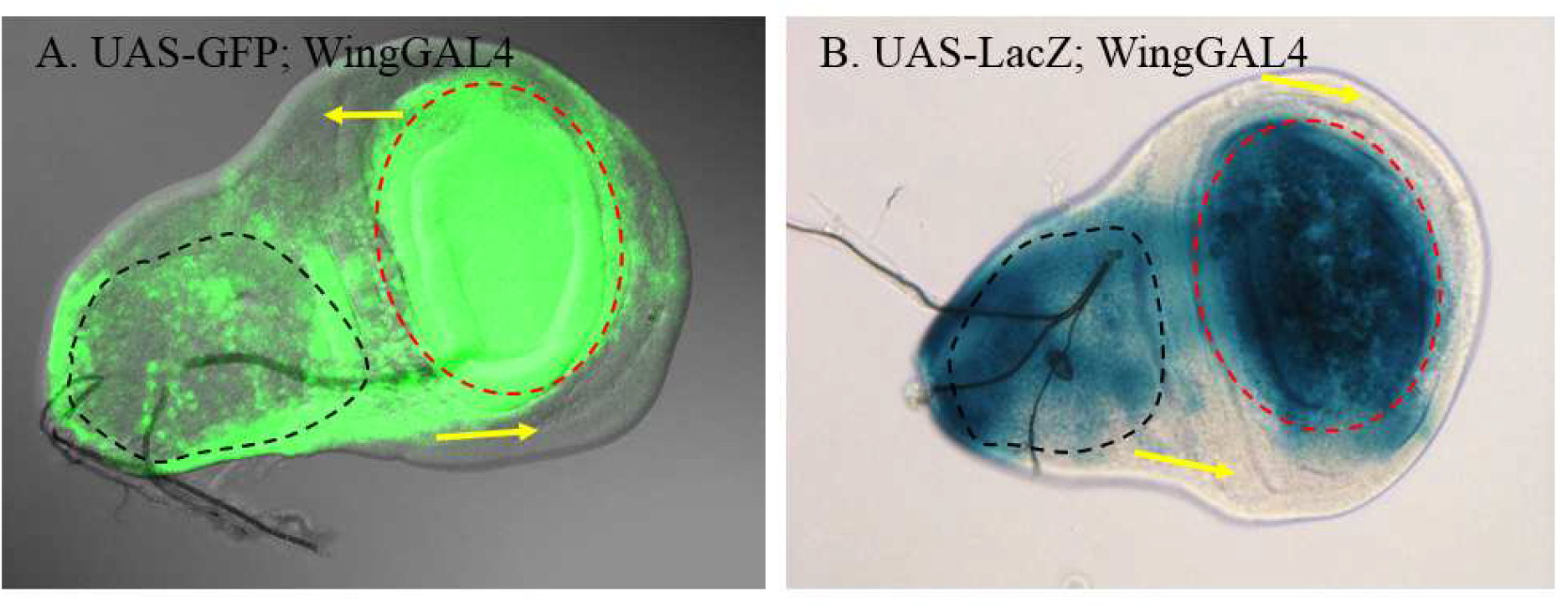
Wing-GAL4 expression pattern analysed by GFP and LacZ staining in wing imaginal discs. A. GFP expresses in the wing pouch (red circle) and notum (black circle) regions of disc, though absent at the margins (yellow arrows). B. LacZ staining shows similar expression pattern.

**Figure S2.**
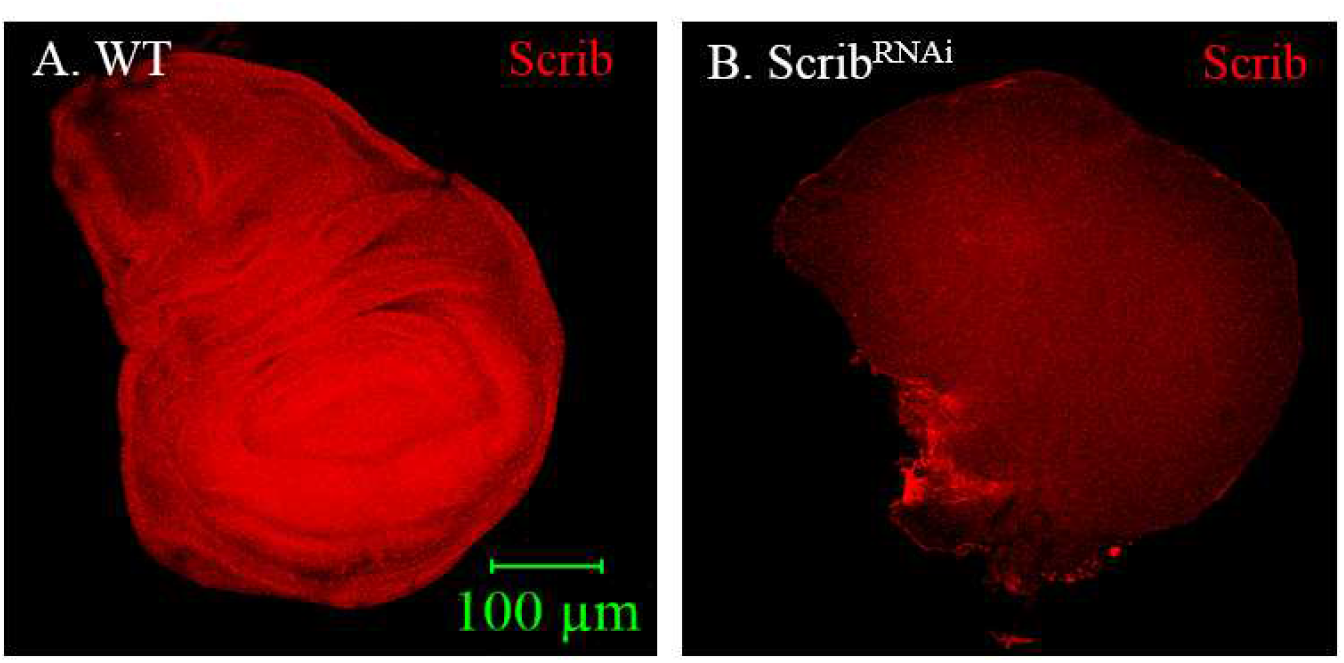
Expression level of Scrib as observed by immunostaining. A, Wild type (WT) shows high level of Scrib (red) compared to B, Scrib knockdown (Scrib^RNAi^) tumorous wing imaginal discs.

**Figure S3.**
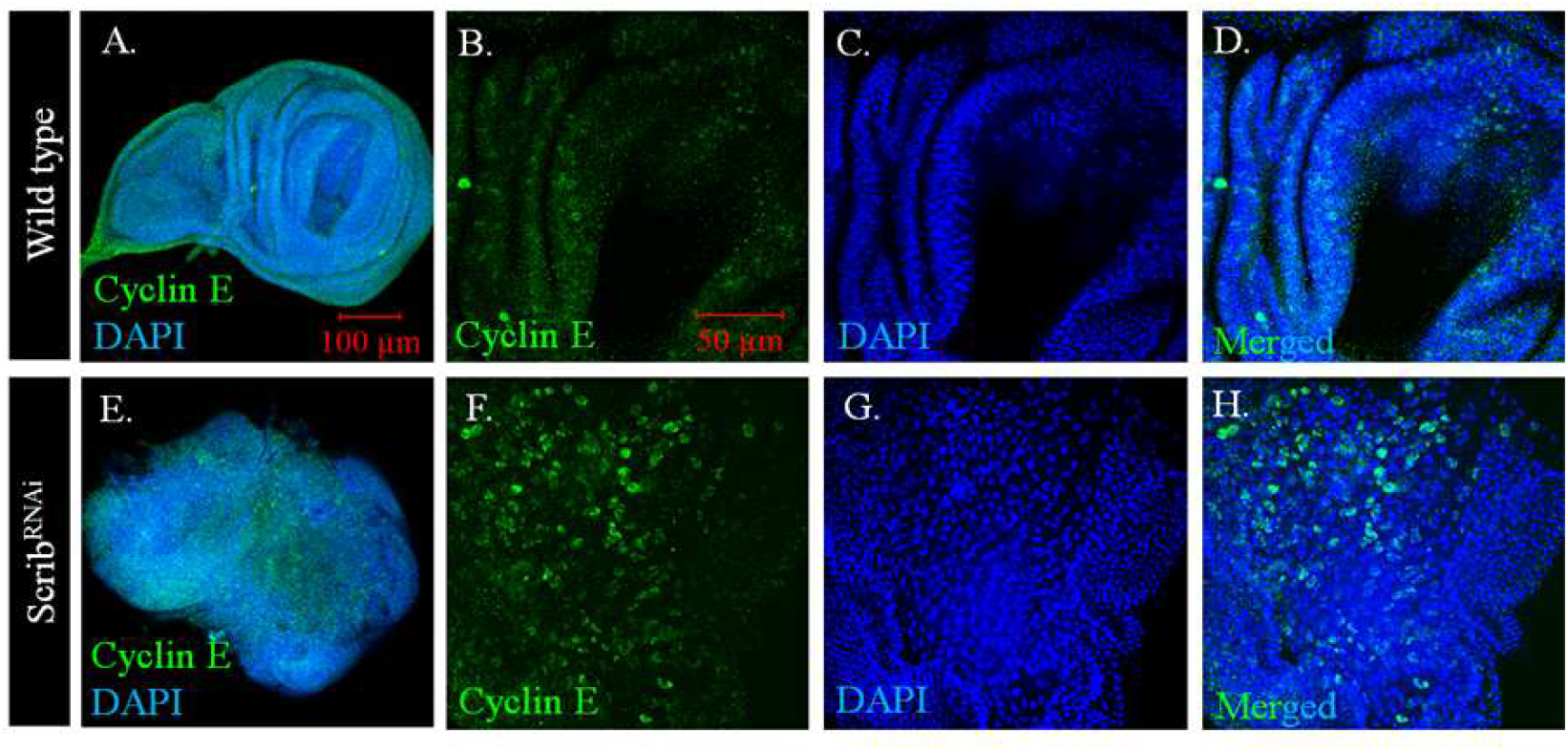
Cyclin-E, a cell proliferation marker, is over expressed in Scrib^RNAi^. Cyclin-E (green) and nuclear staining (blue) in wild type (**A-D**) and Scrib^RNAi^ (**E-F**) wing imaginal discs. **B, F**. Magnified images, cyclin-E expression is higher in Scrib^RNAi^ tumor tissue as compared to wild type. **C** and **G**, nuclear staining with DAPI for wild type and Scrib^RNAi^ respectively. **D** and **H**, merged images for DAPI and Cyclin-E in wild type and Scrib^RNAi^ respectively. Note that Cyclin-E shows nuclear localization.

**Figure S4.**
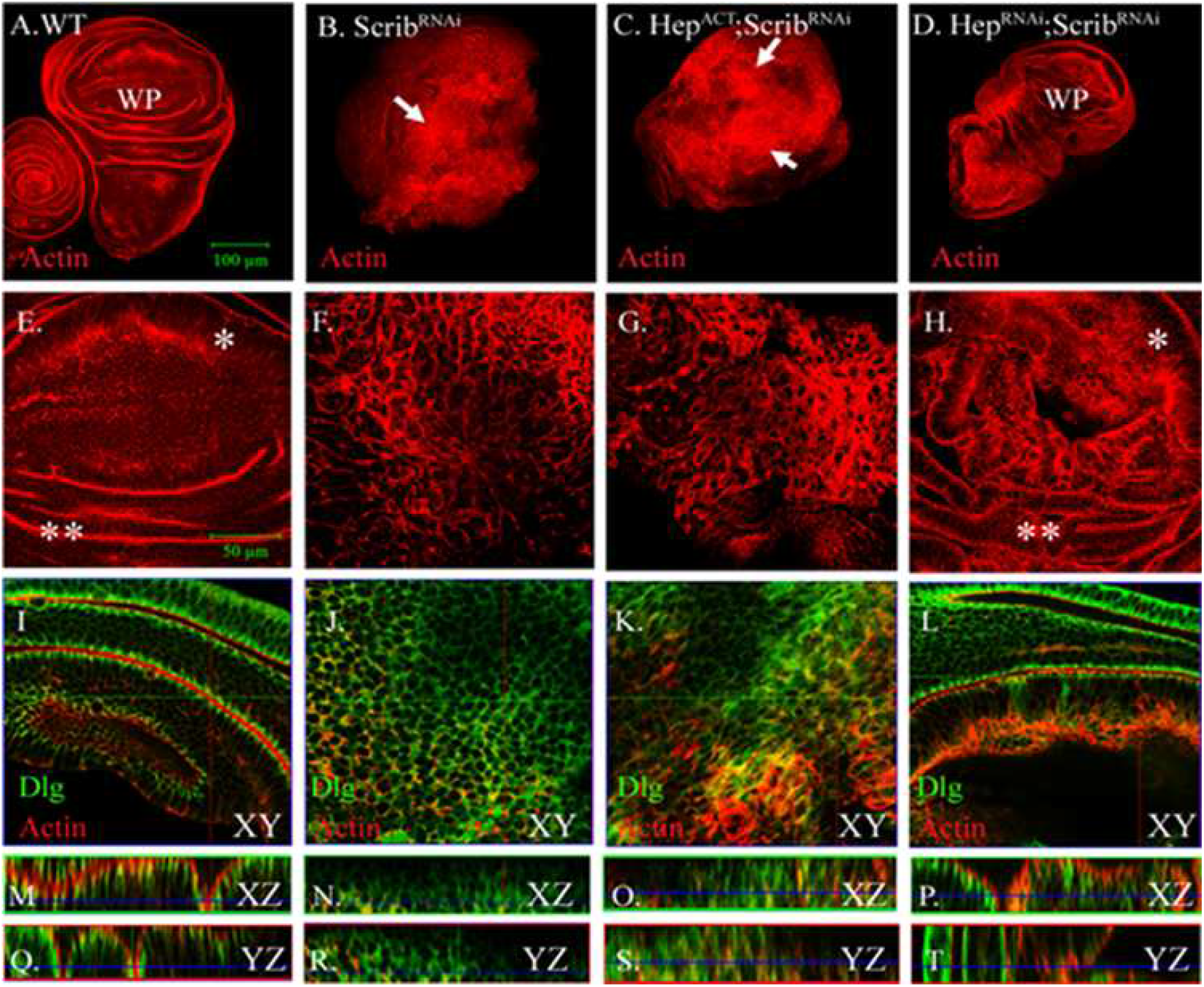
Confocal images of wing imaginal discs representing actin cytoskeleton arrangement and cell polarity. (A) WT wing imaginal disc showing regular actin (red) arrangement and wing pouch (WP) while (B) scrib knockdown causes tumorous growth and distorted actin cytoskeleton arrangement, (C) Constitutive Hep (JNKK) active expression along with scrib knockdown also shows the actin disruption and aggregation while (D) loss of Hep (JNKK) activity by Hep^RNAi^ in scrib knockdown tumor restores actin cytoskeleton arrangement and regular wing structure with wing pouch (WP) formation. (E-H) magnified images represents (E) Wild type wing pouch region with elongated cells (*) and folds (**) while (F-G) shows circular cells and distorted actin arrangement in scrib knockdown and along with active Hep activity respectively while (H) down regulation of Hep activity shows rescue with well-arranged actin cytoskeleton near to wild type. (I-T) represents dlg (green), a baso-lateral cell polarity marker, staining (I, M, Q) Wild type represents regular cell polarity in XY, XZ and YZ plane respectively while (J, N, R) loss of scrib driven tumor shows loss of cell polarity with miss localized Dlg protein. Likewise, (K, O, S) constitutive Hep activated *scrib* knockdown cells also shows the loss of cell polarity, however (L, P, I) reducing Hep activity restores loss of cell polarity in *scrib* knockdown tumor.

**Figure S5.**
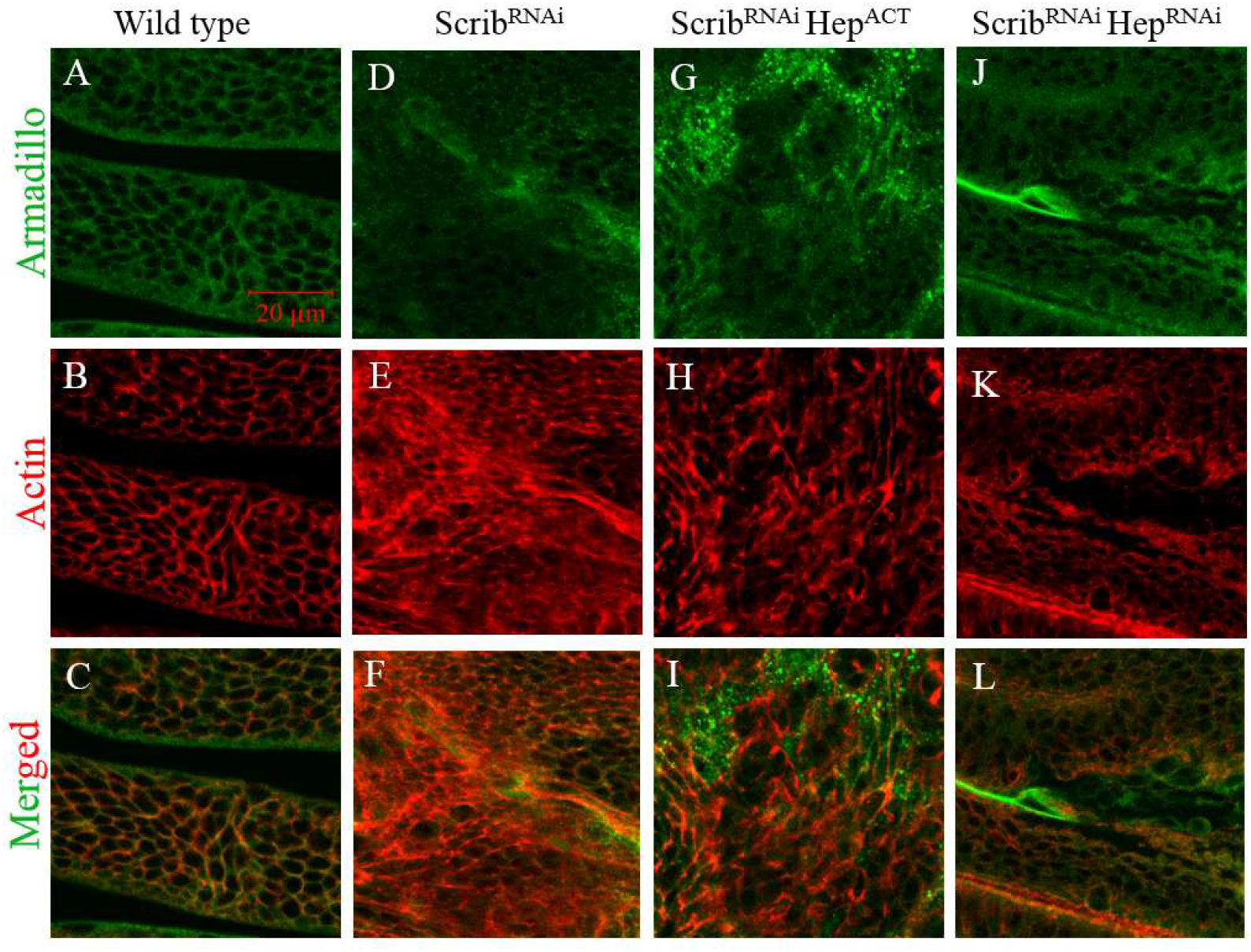
Confocal images of wing imaginal discs representing β-catenin/Armadillo by immunostaining. (A) wild type disc shows armadillo staining (green). (B) phalloidin staining represents actin arrangement in wild type disc (C) merged image of A&B suggests membranous localization of armadillo. While, (D-F) represent armadillo and phalloidin staining in *scrib* knockdown tumor and suggests cytoplasmic presence of armadillo beside membrane. Also, (G-I) constitutive JNK activation (Scrib^RNAi^,Hep^ACT^) enhances cytoplasmic localization of armadillo. While, (J-L) down regulation of JNK signaling (Scrib^RNA^;Hep^RNAi^) restores membranous localization of armadillo.

**Figure. S6.**
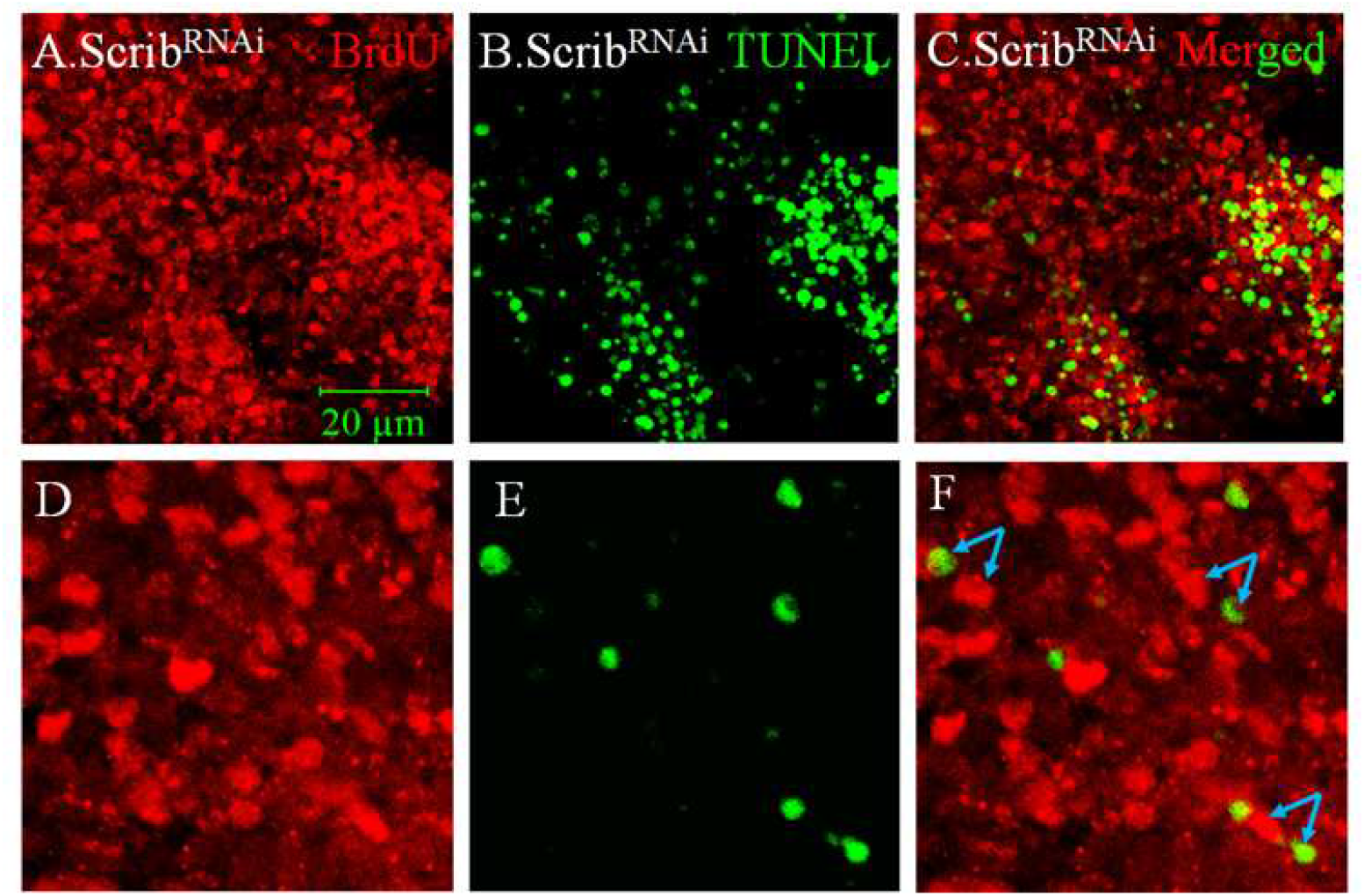
Confocal projection representing BrdU and TUNEL assay in *scrib* abrogated tumor. (A) & (B) represent BrdU (red) and TUNEL (green) positive cells respectively. (C) Shows merged image for BrdU and TUNEL activity. Lower panel magnified images (D-F) show that actively dividing cells are in close proximity to apoptotic cells (blue arrow). It suggests the apoptosis induced cell proliferation in *scrib* abrogated tumor.

